# Vocal motor experiences consolidate the vocal motor circuitry and accelerate future vocal skill development

**DOI:** 10.1101/440388

**Authors:** Michiel Vellema, Mariana Diales Rocha, Sabrina Bascones, Sándor Zsebők, Jes Dreier, Stefan Leitner, Annemie Van der Linden, Jonathan Brewer, Manfred Gahr

## Abstract

Complex motor skills take considerable time and practice to learn. Without continued practice the level of skill performance quickly degrades, posing a problem for the timely utilization of skilled motor responses. Here we quantified the recurring development of vocal motor skills and the accompanying changes in synaptic connectivity in the brain of a songbird, while manipulating skill performance by consecutively administrating and withdrawing testosterone. We demonstrate that a songbird with prior singing experience can significantly accelerate the re-acquisition of vocal performance. We further demonstrate that an increase in vocal performance is accompanied by a pronounced synaptic pruning in the forebrain vocal motor area HVC, a reduction that is not reversed when birds stop singing. These results provide evidence that lasting synaptic changes in the motor circuitry are associated with the savings of motor skills, enabling a rapid recovery of motor performance under environmental time constraints.

## INTRODUCTION

Complex motor skills, such as singing or playing an instrument, are not inherently determined but need to be learned through repetitive practice. Many specialized skills are used only incidentally however, and skill performance degrades during intermittent periods of non-use. This means that skills need to be re-acquired each time the need arises. For example, trained human surgeons need to re-acquire their surgical proficiency after a period of absence (1), food-cashing mammals need to use manipulative skills to retrieve their stored food cashes after periods of hibernation (2), and songbirds need to re-develop high-quality songs to attract a mate after wintering or long-distance migration (3). Whereas learning novel motor skills is generally a lengthy process, the short-term availability of suitable mates and food items imposes a considerable time pressure on the re-acquisition of such specialized skills.

Birdsong, an established model system for vocal motor learning (4), is characterized by a high degree of complex and rapid acoustic modulations. The fine motor skills that are necessary to produce such complex, high quality vocalizations are thought to advertise an individual’s quality through performance-related characteristics such as song complexity and/or production rate (5–8). When juvenile songbirds are one to two months old they start singing noisy, unstructured ‘subsongs’ akin to human babbling (9), which gradually develop into variable, but recognizable species-specific ‘plastic songs’ (10–13). With extensive vocal rehearsal the songs slowly consolidate into high performance ‘crystallized songs’, a process that can take up more than five months in some songbird species (11). For adult songbirds that annually need to re-acquire their songs this process of juvenile song development greatly exceeds the time that is available to attract a mate during the early breeding season. Thus, adult songbirds must considerably speed-up song re-acquisition to quickly produce songs of adequate quality to impress conspecifics. It is currently unclear how songbirds cope with the seasonally imposed time pressure on song development, and what mechanisms may be involved to ensure reproductive success under such time constraints.

The seasonal development and degradation in vocal performance are highly correlated with changes in testosterone levels (14–17), and are often accompanied by gross anatomical and cytoarchitectural restructuring of the brain areas involved in song production (18–24). Thus testosterone-regulated adaptations of the songbird brain may play an important role in optimizing song performance. In this study we used adult female canaries (*Serinus canaria*) to investigate how a learned periodic motor behavior attains a high performance level under developmental time pressure. Although female canaries rarely sing spontaneously, and never produce high quality songs under natural conditions (25), the systemic application of testosterone leads to the development of high-performance song with male-typical features (26–28). This trait provides us with a well-defined behavioral baseline, and allows us to manipulate in detail the onset and offset of song development through repeated testosterone treatments.

Here we demonstrate that during a protracted testosterone treatment, adult female canaries gradually develop stable, species-typical songs through a process of song crystallization similar to what is known for naturally-raised juvenile male canaries (13). Re-treatment with testosterone several months after birds have stopped singing, leads to a rapid recurrence of song performance with song features that strongly resemble those after the first treatment. We propose that once developed, vocal motor memories are retained for an extended period of time, enabling adult animals to recover song performance quickly at a later time, a process termed ‘savings’ (29). We further demonstrate that a subpopulation of neurons in the songbird’s premotor nucleus HVC, a central nucleus in the brain circuitry that controls song production, irreversibly loses a significant number of dendritic spines during the development of vocal skills. This state of reduced spine density is subsequently maintained over long periods in which vocal skills are not used. The observed lasting synaptic pruning could play an important role in the formation of vocal skill savings, enabling birds to rapidly re-acquire song performance within a restricted time period.

## RESULTS

### Testosterone-induced song development in female canaries

To study the development of vocal skills in adult female canaries we recorded and analyzed the entire song ontogeny in acoustically separated animals (5.6 ± 0.6 million song syllables per bird), while manipulating song output by consecutively implanting, removing, and re-implanting birds with testosterone implants.

Whereas no songs were observed prior to treatment, the systemic application of testosterone (T_1_+) stimulated birds to start singing their first songs after three days (3.22 ± 0.46 days). The phases of female song development were similar to those previously reported for male canaries (Fig. 1) (11, 13, 30). The first subsongs were produced in an irregular fashion and consisted of unstructured syllables with few distinctive features. Songs gradually developed into plastic songs in which different phrases of repeated syllables could be clearly distinguished, albeit with variation between syllables of the same type. During the following six months, female canary songs continued to develop, gradually crystallizing into the species-typical stable songs. Although structurally similar to male canary songs, female songs had relatively small syllable repertoires (6.3 ± 1.1 syllable types), consistent with previous reports (28, 31, 32)

**Fig. 1.**
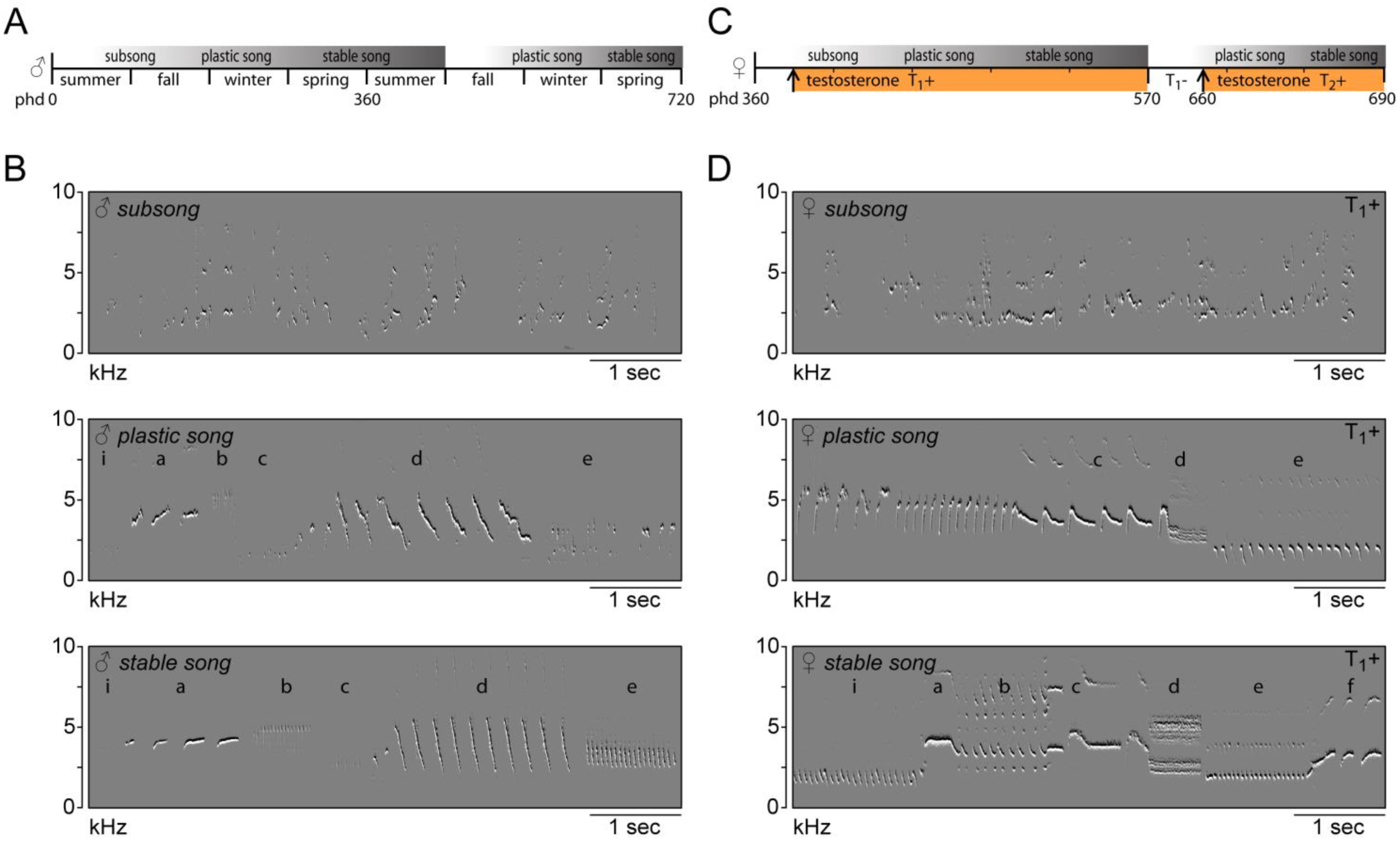
Song in juvenile male and adult female canaries progresses through the same stages of development. **(A)** Schematic of natural song development in juvenile males. **(B)** Example spectrograms illustrating the different developmental stages of male canary song. Subsong was recorded from 45-day old birds, plastic song from 180-day old birds and stable song from 1-year old males. **(C)** Schematic of song development in adult female canaries during a first testosterone treatment (T_1_+), after removal of testosterone (T_1_-), and during a second testosterone treatment (T_2_+). **(D)** Example spectrograms of subsong, plastic song and stable song from an adult, testosterone-treated female canary. Lowercase letters indicate different phrases of repeated syllables.

### Re-application of testosterone triggers an accelerated re-acquisition of song performance

To investigate the birds’ abilities to re-acquire song performance after a period without vocal practice we withdrew testosterone (T_1_-) from the animals, completely abolishing singing behavior in approximately three days (3.14 ± 0.63 days). After 2½ months of no song production birds were treated for a second time with testosterone (T_2_+), inducing song output after three days (3.14 ± 0.74 days). No subsongs were observed after the second testosterone treatment, and all birds were able to produce plastic songs from the first day of singing.

We first compared the development of temporal song features during the 1^st^ and 2^nd^ testosterone treatments (Fig. 2, Fig S1). Particularly the speed at which subsequent syllables are repeated, the syllable repetition rate (SR), has been strongly associated with individual performance (5, 33) and has been shown to increase the attractiveness of the song to conspecifics (34, 35). We observed that during the first two weeks of testosterone-induced female song development, all syllables were typically produced at a similar rate (SR: 10.3 ± 1.3 Hz). While practicing the song, different syllables were gradually sung at different rates, becoming more distinct towards song crystallization (Fig. 2A). Once crystallized, syllables could be distinguished as fast syllables (SR: 21.7 ± 1.0 Hz), medium-speed syllables (SR: 11.9 ± 1.1 Hz), and slow syllables (SR: 4.9 ± 0.7 Hz). A similar distribution of syllable rates re-emerged once songs had stabilized after a 2^nd^ testosterone treatment.

**Fig. 2.**
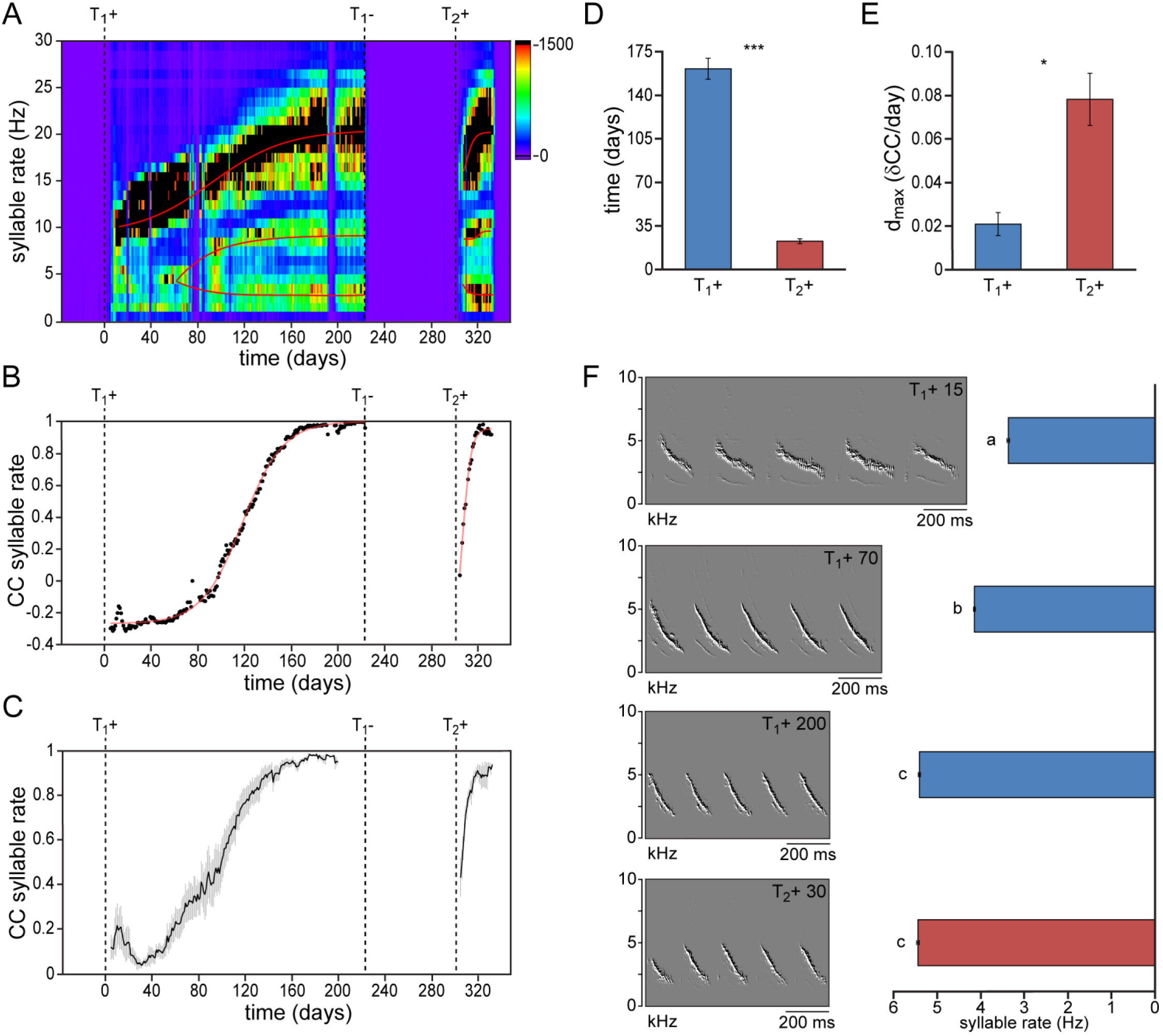
Syllable repetition rates develop faster after prior singing experience. **(A)** Syllable rate histogram from a female canary during two subsequent testosterone treatments. Color scales indicate the number of daily syllables produced for each discrete syllable rate. Curves were fitted through the most occurring syllable rates and are shown in red. **(B)** The correlation coefficient (CC) between the stabilized distribution of syllable rates at the end of the 1^st^ testosterone treatment and each other day during the development and re-development of song. **(C)** The mean syllable rate correlation plot for all animals. Grey bars indicate the SEM. **(D)** Group statistics of all studied birds demonstrating that stable syllable rates were achieved more quickly during a 2^nd^ testosterone treatment (red bars) than during the 1^st^ treatment (blue bars). **(E)** The peak day-to-day increase in the syllable rate CC (d_max_) was higher during a 2^nd^ testosterone treatment than during the 1^st^ treatment. **(F)** Example spectrographs and corresponding bar graphs illustrating syllable rates at 15, 70, and 200 days after a 1^st^ testosterone treatment and 30 days after a 2^nd^ treatment. Columns in D-F represent the mean ± SEM (a,b,c: P ≤ 0.001, ANOVA; * P < 0.05, *** P ≤ 0.001, paired t-test, n = 6).

To determine how fast this syllable rate pattern emerged during subsequent testosterone treatments we took crystallized song from the last week of the first testosterone treatment as a reference pattern, and calculated the correlation coefficient (CC) between this reference and each day of song development and re-development (Fig. 2B,C). While syllable repetition rates gradually developed and stabilized over the course of 160 ± 8 days during the 1^st^ testosterone treatment, stable rates were obtained more than 7 times faster, within 22 ± 2 days, during the 2^nd^ treatment (Fig. 2D; paired t-test: P < 0.001). In addition the maximum daily increase in similarity (d_max_) was also significantly higher during song re-development than during initial song development (Fig. 2E; d_max_: 0.021 ± 0.005 for T_1_+, and 0.078 ± 0.012 for T_2_+; paired t-test: P < 0.05), indicating an accelerated development of song performance in birds with previous singing experience.

Since syllable rate is a compound feature that is determined by both syllable duration and the interval between subsequent syllables, we analyzed syllable durations and pause durations separately. Syllable durations developed slowly during song acquisition, reaching stable values after 153 ± 19 days, while pause durations stabilized in 82 ± 13 days (Fig. S1). Both temporal features re-emerged more quickly during song re-acquisition than during the first song developmental phase (duration: 4.8 ± 1.3 days; paired t-test: P < 0.001; pause: 10.7 ± 2.2 days; paired t-test: P < 0.01; Fig. S1).

To determine the bird’s ability to recover spectral song patterns we analyzed the recurring patterns of frequency modulation (FM), amplitude modulation (AM), bandwidth (BW), mean frequency (MF) and Wiener entropy during song acquisition and re-acquisition. Similar to temporal song features, spectral features were more quickly established during song reacquisition than during the initial song development (Fig. 3, Fig. S2). All observed spectral features demonstrated a gradual development of more than 110 days before reaching stable values. After a period in which the birds did not produce any song, testosterone-induced song reacquisition led to a rapid stabilization of spectral patterns within 16 days (Fig. 3, Fig. S2; paired t test: P < 0.01). Thus both temporal and spectral song features can be acquired significantly faster by experienced birds that have developed singing skills once before.

**Fig. 3.**
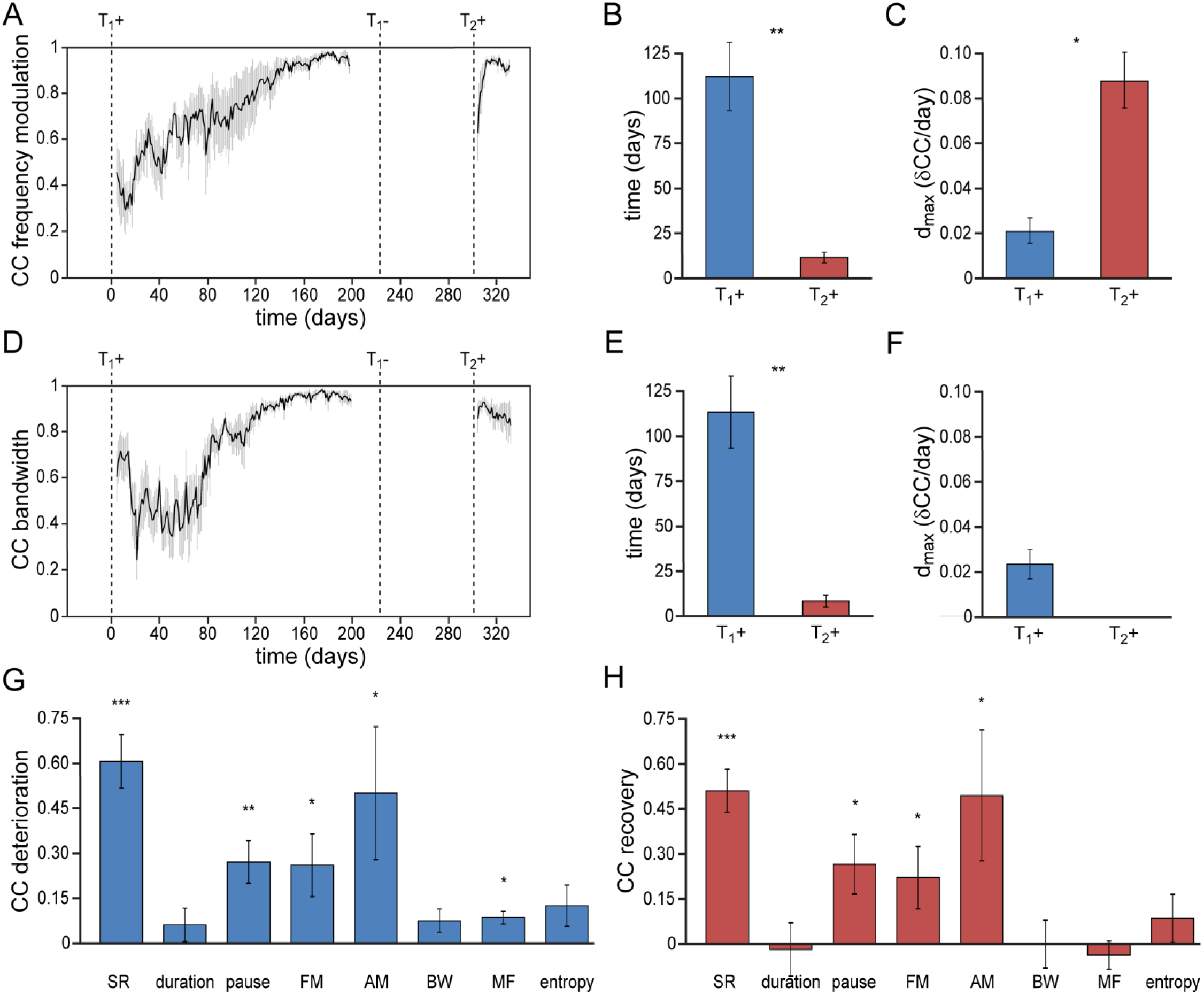
Differential re-development of song features. **(A)** Mean correlation plot for all animals illustrating the development of the frequency modulation (FM) distribution in the song during the development (T_1_+) and redevelopment (T_2_+) of song. The correlation plot illustrates a gradual FM development during T_1_+, followed by a short phase of FM re-development during T_2_+. **(B)** Group statistics demonstrating that the stabilization of the FM distribution in the song took less time during a 2^nd^ testosterone treatment (red bars) than during the 1^st^ treatment (blue bars). **(C)** The peak day-to-day increase in the FM CC (d_max_) was significantly higher during a 2^nd^ testosterone treatment than during the 1^st^ treatment. **(D)** Mean correlation plot for the syllable bandwidth showing a gradual development during a 1^st^ testosterone treatment (T_1_+), followed by an immediate recovery of syllable bandwidth during a 2^nd^ treatment (T_2_+). **(E)** Group statistics demonstrating that stable syllable bandwidths were achieved more quickly during a 2^nd^ testosterone treatment (red bars) than during the 1^st^ treatment (blue bars). **(F)** The peak day-to-day increase in the syllable bandwidth CC (d_max_) for T_1_+. D_max_ could not be calculated for T_2_+, as we observed no developmental increase of this song feature during the 2^nd^ testosterone treatment. **(G)** Deterioration in the distribution patterns of all analyzed song features during absence of song production (T_1_-), and **(H)** subsequent recovery of song features during testosterone-induced re-development of song (T_2_+). Grey bars in A and D indicate the SEM. Columns in B,C and E-H represent the mean ± SEM (* P < 0.05, ** P < 0.01, *** P ≤ 0.001, paired t-test (B,C,E,F) and one-sample t-test (G,H), n = 6).

### The absence of song practice causes selective deterioration of song features

Whereas song re-acquisition in female canaries resulted in a rapid recurrence of stable temporal and spectral patterns, not all song features re-developed in the same way. Most notably, some song features demonstrated a clear deterioration in the absence of vocal practice, while other song features remained stable despite the absence of vocal practice (Fig. 3G,H).

Of the studied temporal song parameters, syllable durations did not show any deterioration in the period that birds did not sing and thus also did not need to be re-acquired (Fig. S1A-C). Both the syllable rate and the pause length between syllables did deteriorate however, and a short phase of re-development was required to recover optimal patterns for both song features (Fig. 2, Fig. S1D-F). The accelerated re-acquisition of syllable repetition rates thus appears to be driven by the need to re-optimize the timing between syllables rather than syllable lengths.

Of the spectral syllable features that were studied, the FM and AM patterns deteriorated when birds stopped singing, but were re-acquired during a short developmental phase after a 2^nd^ testosterone treatment (Fig. 3A-C, Fig. S2A-C). BW, MF, and wiener entropy did not need to be re-acquired, but maintained strong similarities with the initial acoustic patterns despite the absence of song production in the intermittent period (Fig. 3D-F, Fig. S2D-I). Thus song reacquisition after a period without vocal motor practice appears to rely partly on the memorization of previously learned song features and partly on the re-development of lost song characteristics.

### Similarity of song parameters after subsequent testosterone treatments

The song patterns that reappeared during the second testosterone treatment were strikingly similar to those that developed after the first treatment (Fig. 4). Visual inspection of the song spectrograms revealed no differences in syllable repertoire when comparing crystallized songs from the first and second testosterone treatment phases (Fig. 4A,B; 6.3 ± 1.1 syllable types for both treatments). In addition both temporal and spectral features demonstrated a strong similarity in their distribution patterns between subsequent testosterone treatments, with correlation coefficients above 0.82 for all studied song features. No significant differences in the measured correlation coefficients were detected between stable songs from the 1^st^ and 2^nd^ treatment periods for any of the measured song parameters (Fig 4C). These data suggest that independent of the level of deterioration after birds stop singing, testosterone-induced song re-acquisition in canaries is driven towards the previously acquired song pattern.

**Fig. 4.**
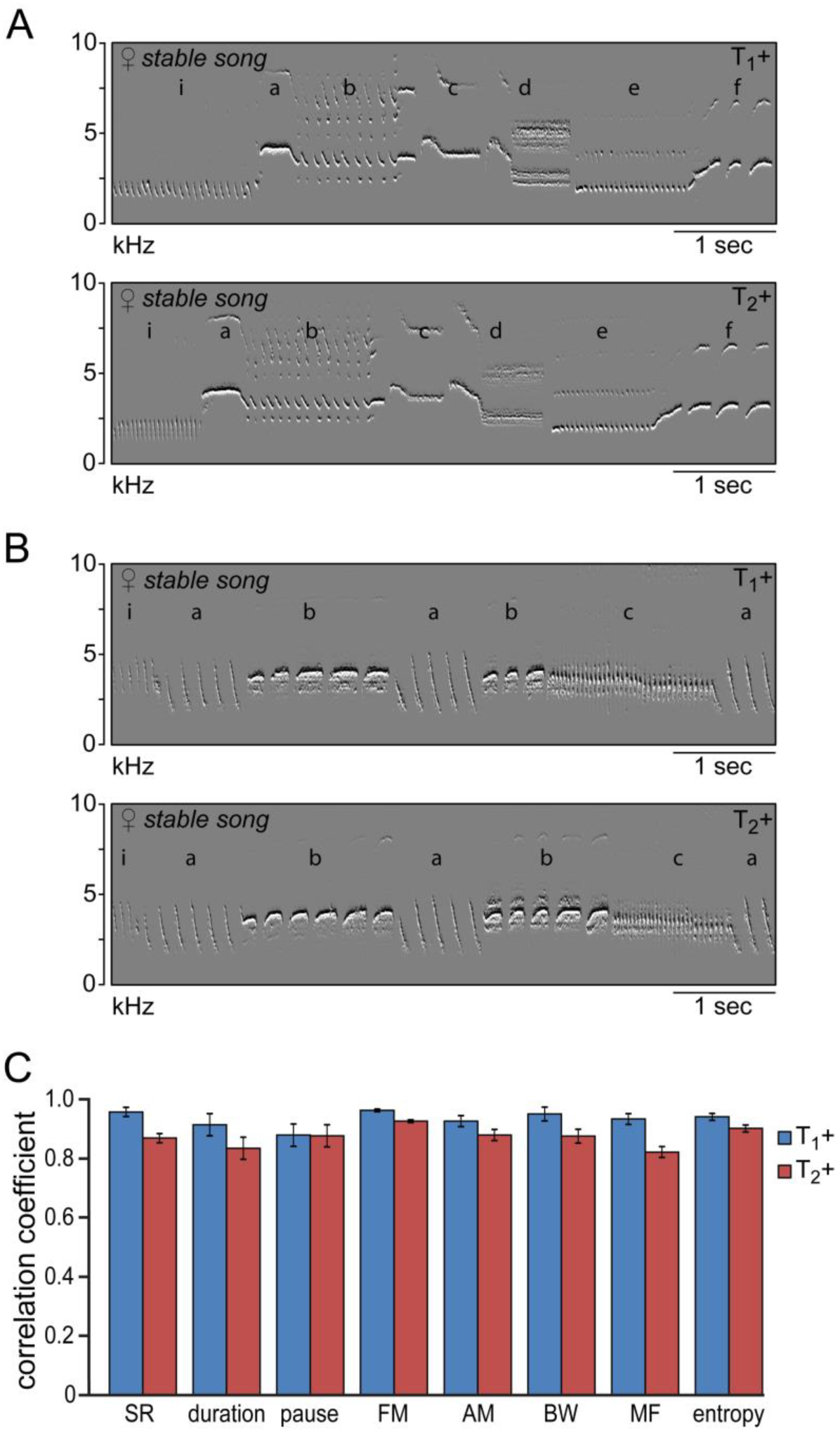
Similarity of time and frequency parameters after subsequent testosterone treatments. **(A,B)** Example spectrograms of stable song from two animals during the 1^st^ (T_1_+) and 2^nd^ (T_2_+) testosterone treatment illustrating the strong similarity in song structure. **(C)** The songs produced during the 1^st^ (T_1_+) and 2^nd^ (T_2_+) testosterone treatment demonstrated a high level of correlation of more than 80% for all analyzed song features. No significant differences where observed within song features between the two developmental phases. Columns in C represent the mean ± SEM (NS, paired t-test, n = 6).

### Singing-related pruning of neuronal dendritic spines

The finding that the re-acquisition of song performance progresses considerably faster than the initial acquisition and leads to the production of highly similar song patterns suggests that singing skills are retained during intermittent silent periods. Motor learning and memory are generally considered to rely on the maturation and consolidation of the synaptic connections within the neural circuitry that drives the behavior in question (36–38). To investigate changes in synaptic connectivity during vocal motor development we quantified the number of neuronal dendritic spines in the forebrain motor nucleus HVC before, during, and after the acquisition of vocal motor skills (Fig. 5).

**Fig. 5.**
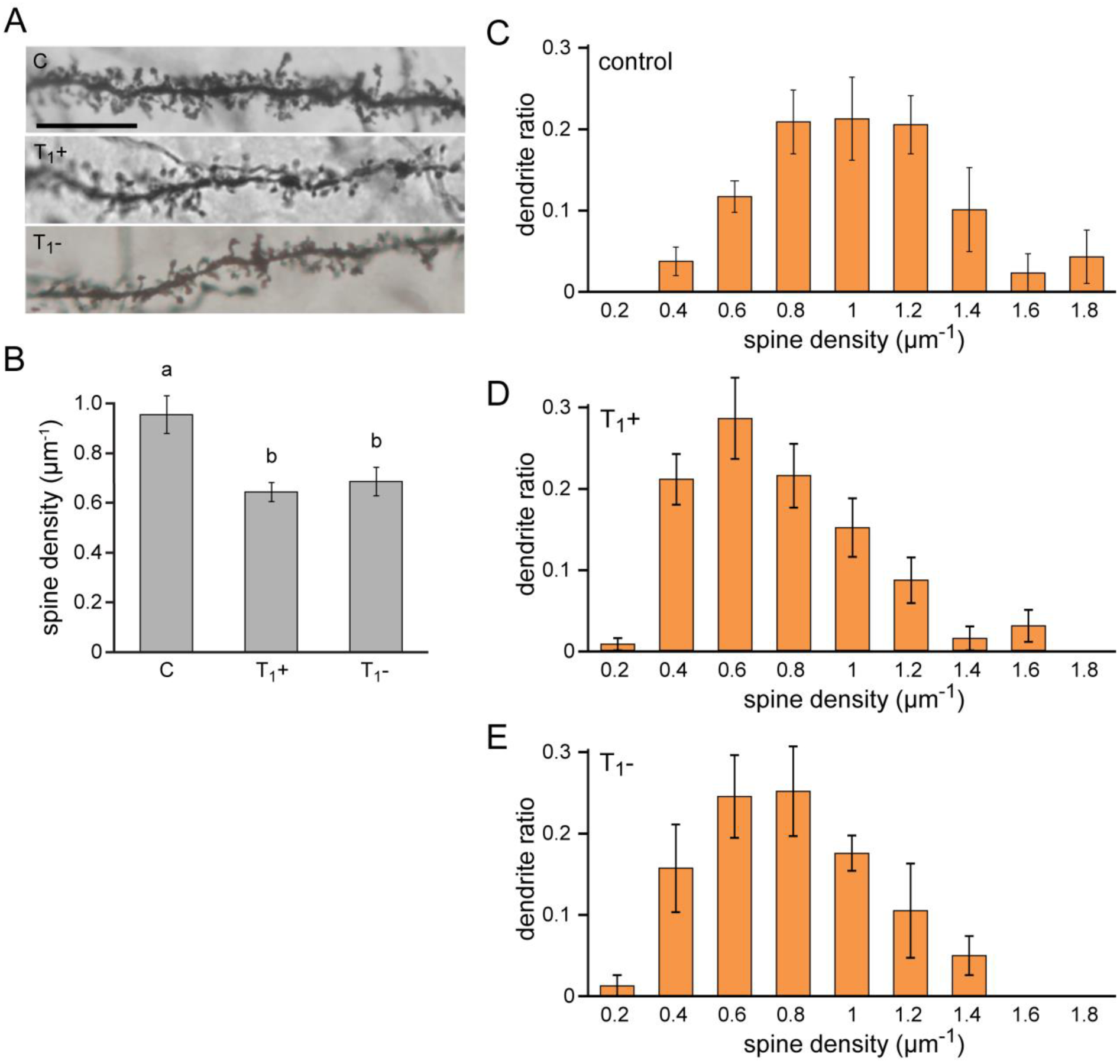
Lasting synaptic pruning in the forebrain motor nucleus HVC. **(A)** Photomicrographs of dendrite segments from non-singing control animals (C), testosterone-treated, singing female canaries (T_1_+), and non-singing individuals with prior singing experience (T_1_-). **(B)** Compared to naive control birds (C), spine densities were significantly reduced in testosterone-treated, singing birds (T_1_+), and remained significantly reduced up to 2.5 months after birds stopped singing by withdrawing testosterone (T_1_-). **(C-E)** The distribution of dendrites based on their spine density demonstrated a shift towards more dendrites with fewer spines in testosterone treated (T_1_+), singing birds and testosterone removed (T_1_-), non-singing birds compared to non-singing control birds. Columns in B-E represent the mean ± SEM (a,b: P < 0.01, ANOVA, n=6). Scale bars = 100 μm.

Excitatory projection neurons in the HVC of testosterone-treated, singing female canaries displayed a more than 30% reduction in dendritic spine density compared to untreated, non-singing control birds (C: 0.95 μm^−1^, T_1_+: 0.64 μm^−1^; Dunnett’s test: P < 0.01), indicating a significant synaptic pruning during the acquisition of singing skills (Fig 5B). Compared to non-singing control birds, spine densities remained significantly lower in birds that stopped singing for 2.5 months after testosterone removal (C: 0.95 μm^−1^, T_1_-: 0.69 μm^−1^; Dunnett’s test: P < 0.01). Thus, the synaptic pruning associated with song acquisition is not reversed when birds stop singing, but instead the pruned state is maintained in periods in which the birds do not sing (Fig. 5B).

Strong differences were observed in the density of dendritic spines between different spinous neurites (range: 0.15 to 2.14 spines/μm). To investigate which neurite types were associated with the observed spine pruning we extracted the distribution pattern of neuronal dendrites in HVC based on the spine density (Fig. 5C-E). The total density of both spinous and aspinous neurites in HVC did not significantly differ between the treatment groups (ANOVA: P > 0.36). Instead of an equal reduction in spines across all neurite types, we observed a shift in the dendrite ratio from densely packed dendrites towards less dense dendrites in the testosterone treatment and removal groups (Fig. 5C-E). Especially in the range of dendrites with more than 0.8 spines per μm a strong reduction was observed in both testosterone-treated, singing birds, and testosterone-removed, non-singing birds compared to non-singing controls (C ratio: 0.64 ± 0.06, T_1_+ ratio: 0.28 ± 0.05, T_1_- ratio: 0.33 ± 0.06; ANOVA: P < 0.01). These data suggest that the observed vocal motor acquisition is accompanied by a lasting synaptic pruning in nucleus HVC, a process that could underlie the memorization and rapid re-acquisition of previously established song patterns.

## DISCUSSION

The ability to learn new skills enables one to adapt to changes in the surrounding environment. Whereas learning new skills requires time and effort, the survival of many species relies on their ability to utilize specialized skills in a timely fashion, capitalizing on the short-term availability of foods, mates, and other environmental factors (2). In this study we establish the repeated, hormone-driven acquisition of vocal motor performance in female canaries as a model system for studying periodically recurring motor behaviors. Using this model, we show that while the development of specialized motor skills can be a lengthy process, motor performance can quickly be re-acquired at a later time, even after extended periods of non-use (Fig. 2 & 3). Such apparent motor savings (29) are biologically relevant for numerous skills in a multitude of species, ranging from periodic foraging skills in dolphins (39), bats (40), and monkeys (41) to courtship displays in crabs (42) and songbirds (43). As the use of these specialized skills is interrupted over longer periods, e.g. during long-distance migration or hibernation (2, 3), such skills need to be reacquired quickly to maximize survivability and reproductive success. Our data suggest that songbirds have the ability to retain previously developed singing skills through a process of selective pruning of the neural circuitry involved (Fig. 5), enabling them to rapidly recover songs of adequate quality to attract a mate before breeding opportunities disappear.

It is not known if adult female canaries can learn songs through sensorimotor integration of an external auditory template as has been demonstrated for juvenile male canaries (44). Our longitudinal data on song development in females does provide the first evidence that at least female songs develop through the same proto-syllabic stages and crystallize in a similar time span as is natural for juvenile males (Fig. 1–3) (11, 13, 30). Considering that the female canaries in our study were acoustically separated during the testosterone-induced development of song suggests that the emerging song patterns are either for a large part intrinsically determined (13, 44), or limited to a previously learned collection of syllable types during juvenile development (45–49). The accelerated song re-acquisition observed in this study indicates that acquiring a motor skill for the first time follows a different developmental trajectory than re-acquiring the same skill at a later time. Whether or not female canaries are able to integrate sensory information in their songs, our data suggest that a combination of procedural learning and memory is at the basis of the rapid re-acquisition of song performance. This is not only evident from the accelerated consolidation of song performance during song re-development (Fig. 2 & 3), but is also supported by the observation that birds can forgo the subsong phase of song development when re-acquiring their songs.

Previous studies on long-term memory in rodents and bats have reported an immediate return of high performance in different food acquisition tasks after hibernation (50, 51) or absence of motor practice (52), without a clear phase of skill re-development. Human visuomotor and eye-blink tasks on the other hand do require a short re-learning phase before previous performance levels are re-acquired (53, 54). Differences in the re-acquisition trajectory between species and skill types could reflect a trade-off between how quickly a skilled response is required versus the ability to modify the skilled response. Hibernators may favor an instantaneous return of foraging skills after hibernation as food intake may pose an immediate requirement for survival. On the other hand, courtship displays, such as singing, may benefit from a short, yearly fine-tuning to optimize the effectiveness of the display. The short phase of vocal practice that we observed during the re-acquisition of vocal motor performance could reflect a songbird’s ability to modify or optimize its song output (11, 13, 17). Whereas male canaries are known to annually reorganize their songs, even when deafened (13), no such modifications were observed in our study. The syllable repertoires of our females were much lower than what is common for males. Such a small vocabulary would limit the possibilities to re-arrange or replace syllables without compromising song complexity, and thus few modifications to female songs can be expected.

Several song parameters were maintained across a period in which birds did not sing, while other song features demonstrated clear deterioration (Fig. 3G,H). Birds were able to recover those deteriorated song features with renewed vocal practice. Song re-optimization thus forms a complex process relying on both memorization of some features and re-development of others in order to obtain the original song pattern. Interestingly, those song features that required a phase of re-development to be optimized can all be associated with constraints in motor performance (5, 55). Online modulation of frequency and amplitude requires active contractile modulation of syringeal muscles (56, 57). Syllable repetition and the inter-syllabic interval also depend on how rapidly syringeal abductor and adductor muscles can open and close the syringeal aperture. Possibly song re-development is limited primarily by peripheral neuromuscular control or th muscle performance itself, and renewed muscle practice may be required to re-optimize vocal performance. Syllable bandwidth is often considered a song feature that is limited by motor constraints (5, 33), but we did not observe deterioration of the bandwidth patterns when singing behavior was absent, nor did we observe any recovery of this feature during song re-acquisition (Fig. 3). Bandwidth would only constitute a performance feature in combination with syllable rate, however. By itself bandwidth is not directly determined by how fast syringeal muscles can contract, unlike syllable rate or frequency modulation. Perhaps the maximum achievable upper and lower frequency boundaries are established during the initial song acquisition, and stored together with other declarative features such as the mean frequency and entropy. If birds can use such declarative features as a base for re-establishing a previously learned song pattern, then only procedural, muscle-dependent song characteristics would need to be re-optimized through vocal practice.

To enable a faster recurrence of a previously learned skill, information from the first learning trajectory must be retained to some extent, and needs to be re-accessible at a later time. The experience-dependent, lasting synaptic pruning observed in the forebrain motor area HVC (Fig. 5) could reflect a structural modification facilitating the long-term memorization of the learned skill, priming the neural motor circuitry for future use of that specific skill. HVC is at a central position between the motor circuitry that controls song output and a forebrain-basal ganglia circuit, analogous to the mammalian cortico-basal ganglia circuit, which feeds back on the motor circuitry for song production (4). Cortico-basal ganglia-like circuits have been strongly implicated in the learning and long-term retention of motor skills in humans (58–62), mammals (63–66), and birds (67–76). Deafening-induced song deterioration in adult zebra finches is further accompanied by a decrease in spine size and stability, without altering spine density, specifically in those HVC neurons that project to the avian basal ganglia, Area X (77). Thus maintaining vocal performance may depend on the structural consolidation of the permanent population of X-projecting HVC neurons (19, 78). In our study we observed a lasting reduction of 50% in the number of densely-spined dendrites in HVC after birds developed song. Previous studies have shown that those HVC neurons that project to Area X predominantly consist of neurons with densely-packed spines (79–85). Thus our observed synaptic pruning may be related to a major reorganization within the HVC to Area X connection, a reorganization that could reflect the long-term memorization of vocal skills. A procedural memory trace within the forebrain-basal ganglia loop could potentially weaken or strengthen synaptic connections within the motor pathway of song production through activity-dependent cellular processes, such as long-term depression or potentiation of synapses (86–90), quickly driving motor patterning to previous levels of performance.

Optical imaging studies have demonstrated a rapid accumulation and stabilization of neuronal dendritic spines in HVC upon developmental song learning in zebra finches (91). Moreover, in the mouse cortex, morphological modifications to dendritic spines have been observed that lasted well beyond the learning experience that caused the spine modifications (52, 92–94). These studies strongly suggest that long-term memory retention of previously acquired skills is facilitated by stably maintained synaptic connections. Motor learning in adult mice is accompanied by a rapid increase in spine formation in the motor cortex, followed shortly after by enhanced, selective spine elimination, returning spine densities to pre-training levels (52, 93). Despite such learning-related spine formation and elimination, the large majority of dendritic spines in the mammalian brain seem stably maintained in adulthood (52, 93, 95–98), contrasting our observation of significant synaptic pruning in adult canaries (Fig. 5).

During juvenile brain development in humans, mammals, and birds, synaptic connections are initially overproduced, followed by protracted, activity-dependent synapse elimination during puberty (95–97, 99, 100). By selectively stabilizing functional synapses while pruning away inactive synapses, brain circuits are thought to reduce neuronal plasticity, consolidating newly learned information and reducing unnecessary redundancy (36). One possibility is that the observed synaptic pruning in the adult canary HVC during song acquisition reflects a developmentally delayed, activity-dependent consolidation of the neural motor circuitry. A recent study in zebra finches has shown that singing prevention during juvenile development can avert developmental spine pruning in the robust motor nucleus of the arcopallium (RA) (101), suggesting that brain maturation can be delayed. Such a developmentally delayed maturation of the song control circuit is consistent with our observation that the delayed, testosterone-induced development of song performance in adult female canaries follows a juvenile male-like developmental trajectory. Thus, in the absence of song production, the motor circuitry that controls song production may remain plastic in adult female canaries, only stabilizing once neural activity within the circuitry increases to drive testosterone-induced song output.

The decision-mechanisms underlying the learning-related, selective stabilization and elimination of dendritic spines remain largely unknown. Sex hormones may play an important role in the stabilization of such synaptic connections during the development of motor skills. Many of the brain areas that are involved in song learning and production express high numbers of androgen receptors (102–106). Furthermore, changes in neuronal dendritic growth and synaptic morphology in the songbird RA have been observed to accompany testosterone-induced song development in female canaries (107–109). In addition, Frankl-Vilches et al. (110) recently observed that 85% of the seasonally and testosterone upregulated genes in the canary HVC are related to neuron differentiation, axon, dendrite and synapse organization. In the brain of birds and mammals, sex hormones are known to upregulate the expression of brain-derived neurotrophic factor (BDNF) (111–113), which in its turn contributes to the stabilization of synapses through the interaction with tyrosine receptor kinase B (TrkB) (114). Furthermore, BDNF has been shown to be essential during testosterone-induced song development in female canaries (32), providing a likely pathway by which sex hormones can consolidate motor skills by stabilizing synapses.

Sex hormones can be regulated by both environmental and social factors (115–117), thereby providing a timed cue for the consolidation of behavioral output and the underlying brain circuitry. Such a cue could facilitate the timely formation and recall of long-term memories of relevant sensory and motor information. Our study suggests that lasting, hormone-driven synaptic pruning of related brain circuitries could form a basis for such long-term memories, enabling a quick recovery of previously acquired skills. The ability to respond quickly to environmental and social cues through such primed hormone-sensitive control mechanisms would constitute an important adaptation to ensure competitive proficiency and maximize reproductive success under environmental restrictions. The results obtained from this study may have implications not only across different animal groups, but may generally apply to different motor, sensory, and motivational systems that rely on a rapid recall of previously learned information.

## MATERIALS AND METHODS

### Subjects

For this study adult domesticated canaries (*Serinus canaria*) were either purchased from a local breeder in Antwerp or taken from the breeding colony of the Max Planck Institute for Ornithology, Seewiesen. All animals were raised and kept under local natural daylight conditions, until the day length reached 11 hours in early March. Birds were subsequently housed individually in sound attenuating chambers for song recordings. Experimental procedures were conducted according to the guidelines of the Federation of European Animal Science Associations (FELASA) and approved by the Ethical Committee on animal experiments of the University of Antwerp.

### Hormone treatment

To stimulate song production in naive female canaries, birds were subcutaneously implanted (T_1_+) with 8 mm silastic tubes (Dow Corning, Midland, MI; ID: 1.47 mm) filled with crystalline testosterone (Sigma-Aldrich Co., St. Louis, MO). During the hormone treatments a light cycle of 11 hours light, and 13 hours darkness was maintained to exclude photoperiodic effects on song production. This light cycle was chosen because under natural conditions it corresponds to a phase of testosterone up-regulation and physiological sensitivity to hormone fluctuations in canaries (14). After consolidation of song we removed the hormone implants (T_1_-) to end song production, and changed the daylight conditions to 8 hours light and 16 hours dark, inducing a molting phase of approximately 1.5 months. Once the molt was finished, the day length was gradually returned to 11 hours over the following month. Subsequently, the song-experienced birds received a second testosterone treatment (T_2_+), and were sacrificed 5 weeks after implantation.

### Hormone analysis

Throughout the experiments, blood samples were collected from the birds’ right wing veins using heparinized haematocrit capillaries (Brand, Wertheim, Germany). Directly after blood collection, samples were centrifuged at 3000 RPM for 10 min to separate cells from plasma, and stored at −80°C for later analysis.

Testosterone concentrations in the blood plasma were determined by radioimmunoassay (RIA) after extraction and partial purification on diatomaceous earth (glycol) columns, following the procedures described previously (118, 119).

Briefly, plasma samples were extracted with dichloromethane (DCM) after overnight equilibration of the plasma with 1500 dpm of ^3^H-labeled testosterone (Perkin-Elmer, Rodgau, Germany). After separation of the organic phase, the extracts were re-suspended in 2 × 250 μl 2% ethylacetate in isooctane and fractioned using columns of diatomaceous earth:propylene glycol:ethylene glycol (6:1.5:1.5, w:v:v). Collected testosterone fractions were dried, resuspended in phosphate-buffered saline with 1% gelatin (PBSG), and left to equilibrate overnight at 4°C. Extraction of plasma testosterone with dichloromethane resulted in 86 ± 11% (mean ± sd) recovery. Hormone samples were incubated with antisera against testosterone (Esoterix Endocrinology, Calabasas Hills, CA), and after 30 minutes ^3^H-labeled steroids (13500 dpm) were added and left to incubate for 20 hours at 4°C. Free steroids were separated from the bound fractions by absorption on dextran-coated charcoal, centrifuged, and decanted into scintillation vials to be counted.

Testosterone concentrations in the plasma samples were measured in one assay. The lower detection limit of the RIA was 0.33 pg/tube, and all measured plasma levels were above the lower detection limit. The intra-assay variation of a chicken plasma pool as control sample at the beginning and end of the assay was 6.7%. Pooled plasma levels of testosterone for all birds during the different consecutive hormone treatments are shown in Table 1. Mean detected hormone levels were within range of physiological plasma levels in nest-building male canaries (range: 360 – 7970 pg/ml; n=6) and previously reported values for other small passerines (120–122).

**Table 1.** Plasma levels of testosterone (T) during different hormone treatments.

| treatment | T levels (pg/ml) | Significance * |
| --- | --- | --- |
| control | 154 ± 45 |  |
| T<sub>1</sub>+ | 7377 ± 770 | P<0.001 |
| T<sub>1</sub>- | 87 ± 24 | NS |
| T<sub>2</sub>+ | 7219 ± 830 | P<0.001 |

\* P-values of comparisons between different hormone treatments against control values (ANOVA with Dunnett's t correction for multiple comparisons against 1 control).

### Song recording

To monitor the entire song ontogeny during consecutive testosterone treatments, naive female canaries (n=6) were kept individually in sound attenuating chambers during testosterone treatment (T_1_+), testosterone removal (T_1_-) and testosterone re-treatment (T_2_+), while song output was continuously recorded. This approach resulted in a song database of 5.6 ± 0.6 million song syllables per bird, recorded over the course of approximately 1 year.

The vocal activity of each bird was recorded using Sound Analysis Pro 2.0 (SAP)(123). Omnidirectional condenser microphones (TC-20; Earthworks, Milford, NH) connected to a multichannel microphone preamplifier (UA-1000; Roland, Los Angeles, CA) were used to acquire and digitize all sounds produced within the sound attenuating boxes with a sampling frequency of 44.1 kHz. The incoming signal was filtered online using an amplitude and Wiener entropy threshold to exclude background noises, and saved in 60 second waveform audio files (16-bit PCM format).

### Song analysis

Vocalizations were segmented into individual syllables with the fully automated Feature Batch module in SAP by applying an amplitude threshold to the sound wave. Because the distance from perch to microphone ranged between 10 and 20 cm only, little variation in recorded amplitude levels can be expected. The amplitude threshold was selected once manually to assure reliable segmentation, and was kept constant during the analyses of all sound phrases within birds.

To filter out non-song vocalizations, syllables were only included when produced within a song bout of at least 750 ms. A song bout was defined as a sequence of sounds traversing the amplitude threshold with an interval of no more than 100 ms. For each syllable, duration (*d*), pause duration (*i*), syllable rate (SR), frequency modulation (FM), amplitude modulation (AM), syllable bandwidth (BW), mean frequency (MF) and Wiener entropy were calculated and stored in MySQL 5.1 tables (Oracle, Redwood Shores, CA). Bandwidth was calculated for each syllable by subtracting the minimum peak frequency from the maximum peak frequency. The syllable rate was defined for each syllable as: *SR_a_ = (d_a_+i_a_)^−1^*, where *d_a_* is the duration of syllable “*a”*, and *i_a_* is the interval between the end of syllable *“a”* and the beginning of the consecutive syllable within the same song bout, irrespective of syllable type.

To investigate the dynamic change of song features over the course of the experiment, daily histograms were obtained from the entire dataset by rounding individual values to the nearest integer, and plotting the number of times those integers occurred each day. To determine the SRs that were most commonly used at the onset and crystallization of song development a non-linear exponential curve (1) was fitted through the peak values of each SR histogram.

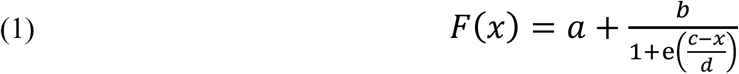

Further developmental correlation plots for all temporal and spectral song features were produced by calculating the Pearson product-moment correlation coefficient (CC) between a 7-day average of the histogram pattern at the time of song stabilization and all other recorded days. This calculation resulted in a value between −1 and 1 for each recorded day, and was used as a measure of similarity where 1 describes an absolute copy of the stable song pattern, and −1 describes the absolute inverse of the stable song pattern. A non-linear exponential curve (1) was fitted through the CC values and song parameters were considered stable as soon as the daily increase in similarity dropped below 0.001. For each song feature we used the fit curve to determine the time from testosterone implant until the feature development stabilized, and to determine the maximum daily increase in similarity (d_max_). Mean CC plots combining all analyzed birds were calculated by averaging the raw (non-fitted) CC values that were calculated for each bird and for each day. We aligned song development in the different birds based on the first day where we detected song.

Spectrographs of the original sound files were visually inspected on syllable-like sound structures to determine the syllable repertoire of each bird and the first occurrence of song after testosterone treatments.

### Specimen preparation

To estimate spine densities and dendrite densities in motor nucleus HVC, brains were collected from a control group of adult female canaries that did not receive any testosterone treatment (C: n=6), a group of birds that was sacrificed five months after testosterone treatment (T_1_+: n=8), and a group of birds that was sacrificed at least 2.5 months after testosterone withdrawal (T_1_-: n=6). All birds were maintained on a light cycle of 11 hours light, and 13 hours darkness and were between 2.5 and 3 years old at the time of sacrifice. Brains were processed for Golgi-Cox staining using the FD Rapid GolgiStain kit (FD NeuroTechnologies, Columbia, MD).

After overnight fixation in a 4% formaldehyde solution in phosphate-buffered saline (PBS; 10 mM; pH 7.4), brains were stored in PBS with 0.05% sodium azide at 4°C until further processing. Brain hemispheres were separated with a razor blade and left hemispheres were immersed for two weeks in an impregnation solution consisting of equal amounts of FD solutions A and B in total darkness at room temperature. Following three days of incubation in FD solution C, brains were cut into 30 μm sagittal sections using a sliding microtome (Leica Microsystems GmbH, Wetzlar, Germany), which were stored in PBS with 0.05% sodium azide at 4°C.

Sections with a regular interval of 180 μm were mounted on SuperFrost glass slides (Menzel GmbH, Brauschweig, Germany), developed in a solution of two parts distilled water, one part FD solutions D, and one part FD solution E for ten minutes, and rinsed in distilled water before embedding in CC/Mount tissue mounting medium (Sigma-Aldrich). Slides were incubated at 70°C until the mounting medium had hardened, cleared in xylene, and further embedded in Roti-Histokitt II (Carl Roth, Karlsruhe, Germany) before coverslipping.

### Spine and dendrite quantification

For each bird, z-stacks of nucleus HVC were obtained from three brain sections with a Nikon Eclipse Ti microscope (Nikon, Tokyo, Japan), equipped with a 60x oil immersion lens (CFI Plan Apo VC 60x Oil). The outline of each HVC section was delineated in ImageJ (http://rsb.info.nih.gov/ij), and 100 μm^2^ non-overlapping regions of interests (ROIs) were randomly placed within the boundaries of HVC prior to the quantifications.

Neuronal dendritic spine densities in HVC were estimated for each brain section by checking ROIs in random order until five ROIs were located that contained spinous dendrite segments (15 ROIs per animal). The individual dendrite segments within the ROIs were traced in ImageJ to measure the length, and the number of visible spines that originated from the traced segment was manually counted. Spine densities were calculated as the number of visible spines per μm of dendrite. All visible spine types were included in the quantifications, including filopodia, long, thin, stubby, mushroom and branched spines (124). For branched spines, each spine head was counted separately.

To further obtain a density measure of spinous dendrites in HVC we counted the total number of spinous dendrite segments in all ROIs examined, and divided the number of observed dendrite segments with the number of ROIs, giving an estimation of the number of spinous dendrites per ROI. Neuronal dendrites were further binned into categories containing 0-0.2, 0.2-0.4, 0.4-0.6, 0.6-0.8, 0.8-1.0, 1.0-1.2, 1.2-1.4, 1.4-1.6, 1.6-1.8 spines per μm. The ratio of neuronal dendrite segments was determined as the likelihood of observing a dendrite segment that falls within one of the bins, relative to the likelihood of observing any spinous dendrite segment. The cut-off value of 0.8 spines per μm was selected as a threshold for densely-packed dendrites, because previous studies have shown that neuronal dendrites with more than 0.8 spine μm^−1^ consist of two dendrite types, the fuzzy dendrites and thick dendrites, while excluding the less densely-spined short dendrites (79, 80).

To obtain a density measure of aspinous neurites in HVC, the number of visible spine-less neurite segments were counted in ten new non-overlapping ROIs that were randomly placed within the boundaries of HVC. We refer to these projections as neurites, because we were unable to distinguish between aspinous dendrites and axons in our tissues.

All quantifications were conducted by an experimenter that was blind to the experimental condition of the animals.

### Statistical analysis

Statistical analyses were performed in SAS 9.3 (SAS Institute, Cary, NC). Differences in spine and dendrite densities between treatment groups were tested with a one-way analysis of variance (ANOVA) with Dunnett’s correction for multiple comparisons against one control value (untreated birds). Changes in hormone levels were determined using a repeated one-way ANOVA with Dunnett’s correction. To detect if the syllable rates of individual syllable types changed over time within the song of each bird, one-way ANOVAs were performed, and when significant, Tukey’s HSD test was used to analyze differences between time points. Song pattern similarities within each bird during the 1^st^ and 2^nd^ hormone treatment were established by calculating Pearson’s CC between an averaged feature pattern histogram from 7 consecutive days of stable song during the 1^st^ treatment with the 7-day pattern average from stable song during the 2^nd^ treatment. Differences between within-individual CCs were compared using paired-samples ttests. Changes in developmental time parameters between subsequent hormone treatments within birds were also tested with paired-samples t-tests. To determine song feature deterioration we calculated the difference between the average CC from the last 7 days of stable song production during the 1^st^ testosterone treatment and the average CC from the first 2 days of song production during the 2^nd^ treatment. Feature recovery was determined by subtracting the average CC from the first 2 days of song production from the average CC from the last 7 days of stable song production during the 2^nd^ testosterone treatment. One-sample t-tests were used to determine if feature deterioration and recovery significantly exceeded zero.

Measurement values in the text are given as means ± SEM, unless stated otherwise.

## ACKNOWLEDGEMENTS

The authors would like to thank Ingrid Schwabl, Monika Trappschuh and Wolfgang Goymann for performing and analyzing the hormone radioimmunoassays. We also thank Sébastien Derégnaucourt for providing male canary songs and Susan Urbanus and Lasse Jacobsen for constructive comments on the manuscript. This work was supported by the Max Planck Society, the European Union’s Horizon 2020 research and innovation program under the Marie Skłodowska-Curie grant agreement nr. 701660 to M.V., the National Research, Development and Innovation Office (NKFIH) grant nr. PD-115730 to S.Z., and the Interuniversity Attraction Poles, grant nr. IUAP-NIMI-P6/38 to A.Vd.L.

## FIGURES

**Fig. S1.**
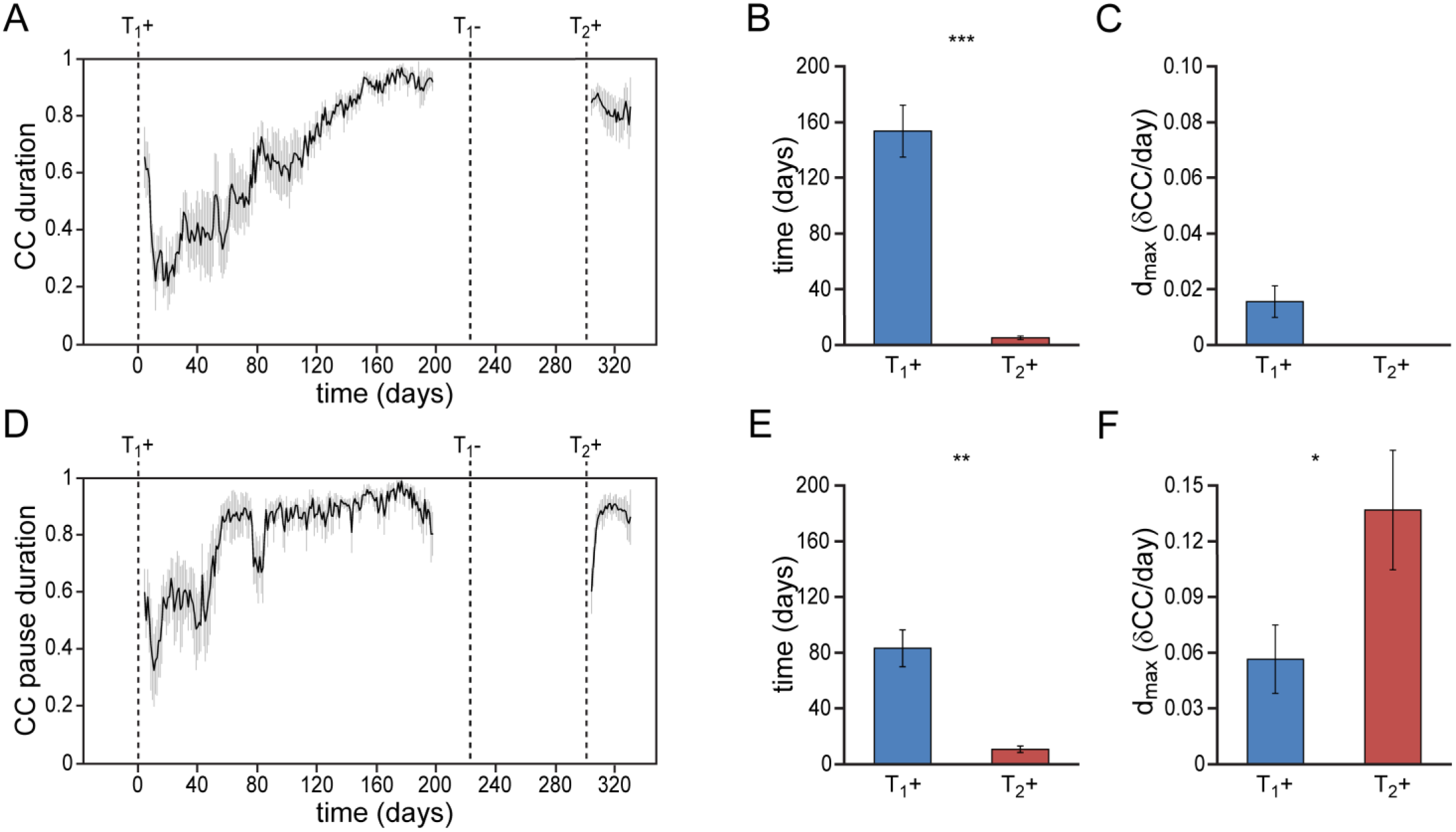
Differential re-development of temporal song features. **(A)** Mean syllable duration correlation plot for all animals showing a gradual development during a 1^st^ testosterone treatment (T_1_+), followed by an immediate recovery of syllable duration during a 2^nd^ treatment (T_2_+). **(B)** Group statistics demonstrating that stable syllable durations were achieved more quickly during a 2^nd^ testosterone treatment (red bars) than during the 1^st^ treatment (blue bars). **(C)** The peak day-to-day increase in the syllable duration CC (d_max_) for T_1_+. D_max_ could not be calculated for T_2_+, as we observed no developmental increase of this song feature during the 2^nd^ testosterone treatment. **(D)** Mean correlation plot for the pause duration between subsequent song syllables. Pause durations stabilized relatively fast during a 1^st^ testosterone treatment (T_1_+), and required a phase of re-development during a 2^nd^ treatment (T_2_+). **(E)** Group statistics demonstrating that stable pause durations were achieved more quickly during a 2^nd^ testosterone treatment (red bars) than during the 1^st^ treatment (blue bars). **(F)** The peak day-to-day increase in the pause duration CC (d_max_) was significantly higher for T_2_+ than for T_1_+. Grey bars in A and D indicate the SEM. Columns in B,C,E,F represent the mean ± SEM (* P < 0.05, ** P < 0.01, *** P ≤ 0.001, paired t-test, n = 6).

**Fig. S2.**
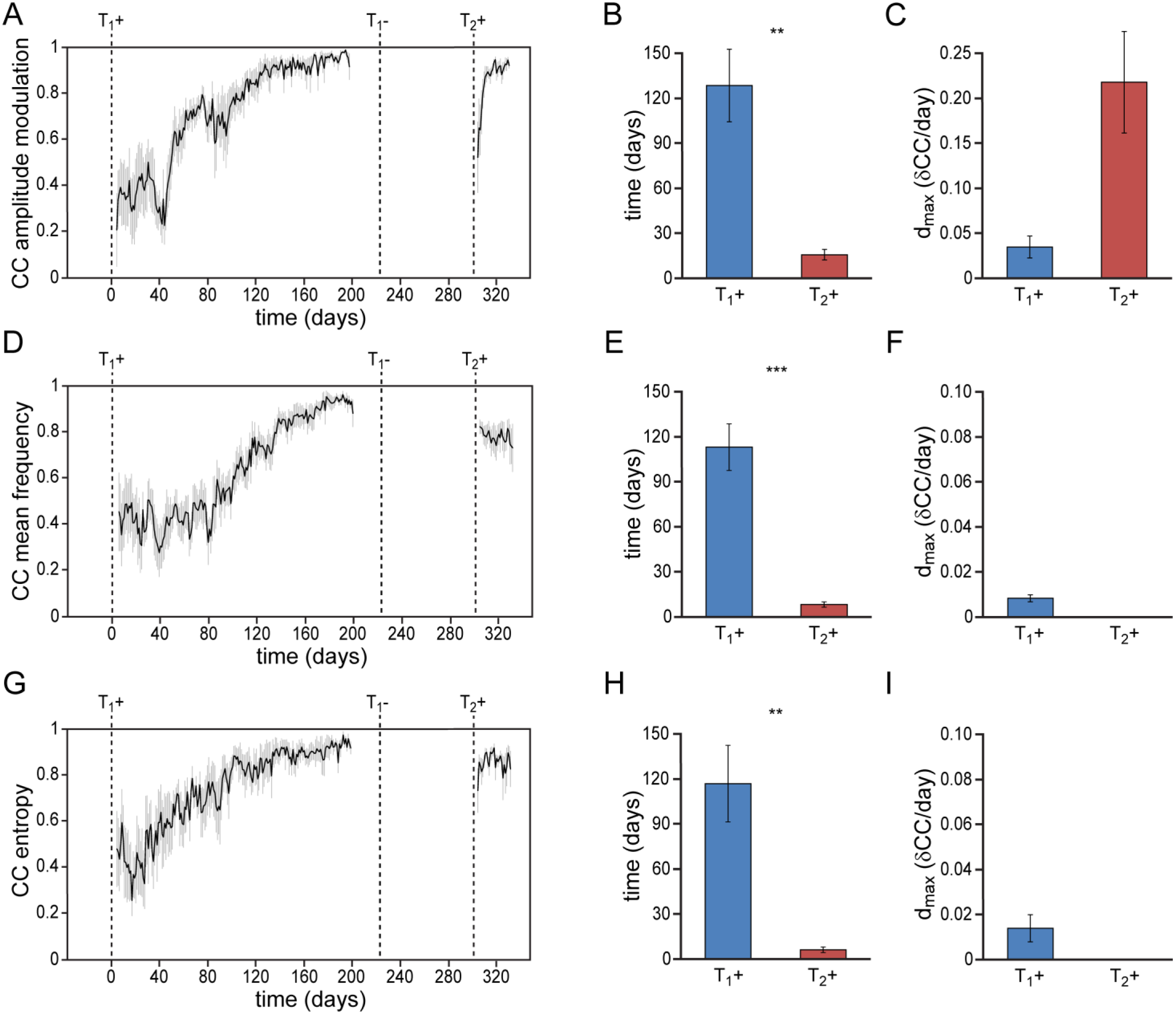
Differential re-development of spectral song features. **(A)** Mean amplitude modulation (AM) correlation plot for all animals. The AM distribution demonstrated a gradual consolidation during the 1^st^ testosterone treatment (T_1_+), and displayed a short phase of re-development during a 2^nd^ testosterone treatment (T_2_+) **(B,C)**. Both mean frequency **(D-F)** and wiener entropy **(G-I)** demonstrated a gradual development during the 1^st^ testosterone treatment (T_1_+) followed by an immediate recovery during a 2^nd^ treatment (T_2_+). D_max_ could not be calculated for T_2_+ in F,I, as we observed no developmental increase of this song feature during the 2^nd^ testosterone treatment. Grey bars in A, D and G indicate the SEM. Columns in B,C, E,F, and H,I represent the mean ± SEM (** P < 0.01, *** P ≤ 0.001, paired t-test, n = 6).

